# Scoring thermal limits in small insects using open-source, computer assisted motion detection

**DOI:** 10.1101/2022.12.20.521307

**Authors:** Fernan R Perez-Galvez, Annabelle C Wilson, Sophia Zhou, David N Awde, Nicholas M Teets

## Abstract

Scoring large amounts of thermal tolerance traits live or with recorded video can be time consuming and susceptible to investigator bias, and as with many physiological measurements, there can be trade-offs between accuracy and throughput. Recent studies show that particle tracking is a viable alternative to manually scoring videos, although it may not detect subtle movements, and many of the software options are proprietary and costly. In this study, we present a novel strategy for automated scoring of thermal tolerance videos by inferring motor activity with motion detection using an open-source Python command line application called DIME (Detector of Insect Motion Endpoint). We apply our strategy to both dynamic and static thermal tolerance assays, and our results indicate that DIME can accurately measure thermal acclimation responses, generally agrees with visual estimates of thermal limits, and can significantly increase the throughput over manual methods.

**Summary statement:** Motion detection algorithm for reliable, automatic scoring of thermal limits in insects with open-source tool

## Introduction

Temperature influences nearly every aspect of an ectotherm’s biology, which has fueled the measurement of thermal limits in a variety of organisms (Dallas and Rivers-Moore, 2012; Lutterschmidt and Hutchison, 1997b). The thermal tolerance of an ectotherm is often conceptualized with a thermal performance curve, which describes the effects of temperature on a metric of biological performance (Schulte et al., 2011). Thermal limits are the minimum and maximum temperature at which a biological process can occur, and motor performance is perhaps the most used metric for assessing thermal limits. Scoring thermal limits involves increasing or decreasing the temperature until motor activity ceases. Alternatively, insects can be exposed to a static, extreme thermal condition until motor failure occurs, and recent work indicates that dynamic and static thermal tolerance measures are mathematically, and perhaps physiologically, related (Jørgensen et al., 2019; Rezende et al., 2014). Further, thermal tolerance can be assessed by measuring the resumption of activity after a period of paralysis, as is the case for the commonly used chill coma recovery time (Sinclair et al., 2015). In insects, thermal tolerance provides physiological information directly related to fitness and is relevant for a number of research areas, ranging from basic ecophysiology (Lee Jr et al., 2006) to the impacts of climate change on insect diversity (Garcia-Robledo et al., 2016).

When an ectotherm approaches its thermal limits, it begins to be physiologically and behaviorally impaired. For critical thermal maxima (CT_max_) or heat knockdown time (HKDT), the sequence of responses includes the loss of righting response, the sudden onset of muscular spasms, and finally the cessation of movement interpreted as “heat rigor”, “coma” or “death” (Lutterschmidt and Hutchison, 1997b). The original description of upper thermal limit endpoints by Hutchison (1961) as the temperature “at which locomotory activity becomes disorganized and the animal loses its ability to escape from conditions that will promptly lead to its death” allows multiple interpretations. While the onset of muscular spasms was an endpoint considered physiologically comparable across phyla (Lutterschmidt and Hutchison, 1997a), the precise criteria can vary, making it difficult to compare results across studies (Sponsler and Appel, 1991). During chilling, a similar series of events occurs, where an ectotherm first slows or stops its normal activity, followed by the loss of coordination that impedes locomotion (i.e., the critical thermal minimum [CT_min_]), and finally, at lower temperatures, movement ceases altogether (i.e., chill coma) (Hazell and Bale, 2011). However, in practice, typically only the CT_min_ is reported and is often assessed by recording failure of a locomotor behavior (typically righting response, ability to cling to a surface, or a response to stimulus) (Sinclair et al., 2015). In contrast, scoring of chill coma recovery time (CCRT) has been typically interpreted as the moment when the insect is “able to stand on its legs”(David et al., 1998). Thus, there are many options available for assessing thermal tolerance, and clear, consistently applied endpoints are paramount for precision and repeatability.

In recent years, there has been an increase in large phenotypic screens to compare thermal tolerance across species (Kellermann et al., 2012; MacLean et al., 2019) or across genotypes of the same species (Gerken et al., 2015; Lecheta et al., 2020; Ørsted et al., 2018). Scoring thermal tolerance traits in real-time or with recorded videos is time consuming, and there can be trade-offs between accuracy and throughput. Thus, methods for automated scoring of thermal tolerance are necessary for improving repeatability and reducing strain on investigators. Recent work has applied particle tracking software (i.e., EthoVision XT) to investigate thermal tolerance in insects, by estimating the onset of coma or recovery using subject displacement, summarized in time intervals, as a proxy for movement (Laursen et al., 2021; MacLean et al., 2022). In Laursen et al. (2021), discrepancies between scored estimates were attributed to the lack of sensitivity of distance-based methods to detect subtle movements of appendages. In MacLean et al. (2022), estimates derived using speed recapitulate the expected biological trends between treatments but present numerical discrepancies against those scored manually. Thus, while particle tracking appears to be a viable alternative to manually scoring videos, it may not detect subtle movements characteristic of some insects, and many of the software options (e.g., EthoVision XT) are proprietary and costly.

In this study, we present a novel strategy for automated scoring of thermal tolerance videos by inferring motor activity with motion detection. The strategy includes two steps: first, video recordings of thermal performance are transformed to a numerical vector of motion, and second, thermal limits are identified from the numerical vector using a scoring method. Here, we test three scoring methods and compare their results against estimates obtained visually to identify a computational interpretation of thermal limits that is most reliable. Our strategy is flexible, and we apply it to both dynamic (CT_max_ and CT_min_) and static (HKDT and CCRT) assays. Our method can accurately measure thermal acclimation responses, generally agrees with visual estimates of thermal limits, and can significantly increase the throughput over manual methods. We provide an open-source Python command line application we call DIME (Detector of Insect Motion Endpoint) that can be used to transform videos to motion data and apply our preferred scoring method MF (Median Frame).

## Methods

### Preparation for thermal performance assays

Two dynamic assays (CT_max_, CT_min_) and two static assays (HKDT, CCRT) of thermal tolerance were conducted in the Oregon R strain of *Drosophila melanogaster* Meigen. Each thermal tolerance assay was performed three times in blocks of 30 flies including five males and five females per acclimation treatment resulting in a total of 30 flies per treatment per assay. Fruit flies were reared in cornmeal-yeast-molasses diet at 25°C in a 12:12 L:D cycle. After emergence, experimental subjects were exposed to one of three acclimation treatments (18, 25, and 30°C) for a period of five days in programmable incubators (MIR-154, Panasonic Healthcare Co., Ltd., Japan) in a 12:12 L:D cycle. On the sixth day, flies were transferred to individual wells in custom acrylic observation arenas using aspirators without anesthesia. One side of the arena was sealed with a transparent acrylic lid to facilitate recording, while the other side was sealed with nylon mesh to allow gas exchange. Flies that were injured during the loading process were removed from the analysis.

Observation arenas containing flies were placed in the center of a programable incubator to perform thermal tolerance assays, which were recorded using a webcam (Logitech V-U0028). In the case of the dynamic assays, CT_min_ and CT_max_, a 0.25°C/min cooling or heating ramp starting at 25°C was programmed in the incubator with a function that changes temperature 2.5°C every 10-minute interval. Even though the program includes discrete temperature steps, the cooling capacity of the incubator, coupled with thermal buffering by the arena, led to an approximately linear thermal ramp (R^2^ CT_min_: 0.999, 0.998, 0.999; R^2^ CT_max_: 0.999, 0.999, 0.986), as measured in the well microenvironment with a DHT22 temperature sensor (Aosong Electronics Co., LTD, China) and an Arduino Nano microcontroller platform (Arduino SRL, Italy). After the experiment, a linear model was fit to the cooling or heating section of the ramp and the linear coefficients were used to translate the time of knockdown into a CT_min_ and CT_max_. The slopes measured in both cases were constant along the duration of the assay but slightly less steep than programmed (CT_min_: -0.227, -0.223, -0.230°C/min; CT_max_: 0.236, 0.238, 0.230°C/min). In static assays, a constant temperature of 36.5°C was used in HKDT, and for CCRT chill coma was induced for 2 hours at 0°C and recovery at 25°C.

### Step 1: Transformation of insect motor activity

Our command line application DIME transformed biological activity in thermal performance videos to a numerical variable of motion detection. DIME was developed in Python v3.8.8 and makes use of the computer vision library OpenCV v4.5.3 (Bradski, 2000). The numerical variable is a measurement of relative pixel intensity change (rPIC) within a region-of-interest (ROI). Ideally, each ROI contains a single individual on a constant background. A video is analyzed as a series of images representing single frames of video data (videoframe) which are extracted as pixel matrices in the red-green-blue (RGB) color space and transformed to a single grayscale value using the standard-definition luminance formula Y’ = 0.299R + 0.587G + 0.114B as implemented in the OpenCV command `COLOR_RGB2GRAY`. The procedure continues by computing the difference between pairs of consecutive grayscale pixel matrices to generate a series of difference matrices. Motion between videoframes is encoded on each pixel of the difference matrix as a deviation from 0. Two filters are applied to each difference matrix, a Gaussian blur to remove flickering particles in the background using the OpenCV command `GaussianBlur` (vertical and horizonal kernel size [*k*] = 7) and a dilation filter to maximize the difference between areas of change with the command `dilate` (convolution iterations [*i*] = 3). Finally, motion detection per individual is achieved by calculating the average absolute pixel intensity change per ROI in the filtered difference matrix.

### Step 2: Inference of thermal tolerance endpoints

Three computational methods were used to interpret the detected motion data: Arbitrary Threshold (AT), Median Frame (MF) and Change Point (CP). The AT method scores a thermal limit as the last or first event of motion at or above an rPIC threshold that was empirically determined by comparing visual estimations with automated estimations (see analysis below). This threshold requires calibration per species per video setup. For each assay, we inferred an optimal threshold in a subset of the data by calculating the x-intercept of a fitted line of measurement error (difference between AT and visual estimation) versus the threshold values used to score AT (Fig. S1). Method MF applies the same rationale to score an endpoint but automatically defines the scoring threshold with a central measure of tendency. Specifically, MF method scores a motor activity endpoint as either the last or first motion event detected equal or above the median of all non-zero rPIC values per individual. The third method, Change Point (CP), statistically identifies the point where an individual changes from being active to inactive, or vice versa. In this interpretation, the method CP considers motor activity in rPIC as a sequence of observations with an underlying pattern where the initial and final means are different, and the change-point between them is unknown (Hinkley, 1970). Here, we apply a maximum likelihood estimator to identify a single change-point of motion along the thermal performance assay of each experimental subject using the “at most one change” option as implemented in the R library `changepoint` v2.2.3 (Killick and Eckley, 2014).

### Step 3: Analysis of computational scoring reliability

Each experimental block was scored visually by one of the authors (block1 1: FRPG, block 2: AW, block 3: SZ) who had different experience scoring thermal limits to simulate a realistic large-screening experimental setup. All statistical analyses were conducted using R v4.1.0 (R Core Team, 2021). First, an exploratory analysis of variance (ANOVA) was conducted in joint datasets containing the visual and computational estimates to identify variance associated to methodology, using the model:

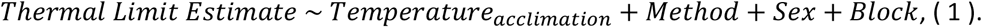

A post hoc Tukey test for honestly significant differences (HSD) was applied to identify significant average differences between computational and visual methodologies. No significant variance associated to the variable *Sex* was detected in lower thermal tolerance assays (CT_min_ and CCRT), so we decided to exclude this variable from the rest of analyses. Also, significant variance associated with the variable *Block* was observed. With this information, we fitted datasets from independent scoring methods to a mixed effects model using *Temperature*_*acclimation*_ as fixed effect and *Block* as random effect with the R library `lmer` from the `lme4` v1.1-29 package (Bates et al., 2015), using the equation:

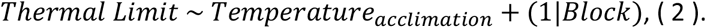

Population marginal means, their associated standard errors, and the post hoc Tukey HSD test to pairwise differences between treatment levels were calculated using the package `emmeans` v1.8.0 (Lenth, 2022). The Kenward-Roger approximation of degrees of freedom was applied when testing independent scoring methods to account for small and unbalanced datasets (e.g., when a methodology was not able to provide an estimate).

Inter-method agreeability between computational and visual estimates was measured with the concordance correlation coefficient (CCC), which evaluates the degree to which pairs of measurements fall in the 45° line of perfect correlation (Lawrence and Lin, 1989). In addition, CCC can provide information on the source of disagreement when decomposed into the bias corrector factor C_b_, a measure of accurateness, and the Pearson correlation coefficient ρ, a measure of precision. Accuracy in C_b_ measures how far the best fit line deviates from the 45° line; ρ measures how far each observation deviated from the best fit line. In our case, CCC and their components were computed as implemented in the R package `DescTools` v0.99.45 (Signorell et al., 2019) in paired datasets containing one computational (AT, CP, MF) and the visual dataset per thermal tolerance assay. Finally, individuals presenting outlying scoring differences were identified with an agreement test (Martin Bland and Altman, 1986) to investigate the cause of disagreement. For each thermal tolerance assay, we calculated the mean difference 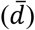 and the standard deviation of the difference (*s*) between visual and MF datasets to estimate the “limits of agreement” at 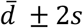.

## Results and discussion

### Scoring efficacy by computational methods

Motor activity of every experimental subject was transformed into a sequence of motion events with our computational tool DIME. Average processing time including transformation and scoring was 18 minutes (1.80 GHz processor) per 2.5-hour long video (640×360 pixel) in contrast with 1 hour spent inspecting the same 30 subjects visually. Videoframe manipulation spends most of the computing resources, as processing time doubles with resolution. Greater throughput can be achieved by increasing the number of subjects per video and increased processing power.

The first two computational methods (AT and MF) are heuristics designed to recapitulate the strategy we currently use to score: scrolling the complete video in reverse (CT_max_, CT_min_, and HKDT), or forward (CCRT), until the endpoint of motor activity (onset or termination) is identified and used to score an estimate. The visually informed method AT provided estimates for every individual tested, with the exception of CCRT where three individuals presented motion activity below the optimal threshold defined. The automated method MF was the most reliable in terms of scoring efficacy as it provided estimates for every individual in all four thermal limit assays. The method CP scored most individuals in upper thermal limits but failed to provide a meaningful score for CT_min_ and CCRT, which could be influenced by the reduced amount of motion information present in subjects exposed to lower temperatures. For this reason, only upper thermal limits obtained by CP method were considered for the following analysis.

### Variance introduced by methodology and recapitulation of treatment effects

In a linear model to estimate sources of variation in thermal limits, the variable *Method* contributed significant variance in CT_max_, HKDT, and CCRT but not in CT_min_ (Table S1), suggesting that the introduction of variance is assay specific. CT_max_ presented the greatest differences between automated and visual scoring methods (Fig. 1). Both AT and MF resulted in statistically significant deviations from visual estimations (−0.27 and -0.39°C, respectively), whilst on a lesser magnitude than CP which deviated by -1.3°C. Despite the differences, computational methodologies consistently recapitulated treatment effects in every case (Table 1). In the case of CT_min_, no average differences were observed between computational and visual estimates and the same aggrupation was recapitulated in every case. On the other hand, static assays (HKDT and CCRT) presented greater variation between scoring methodologies. On average, HKDT estimates obtained by AT and MF methods are not statistically different from those scored visually, but CP estimates differ by -15.9 min. Interestingly, a biological trend was recapitulated by MF and CP methods, but not by those scored visually or visually informed (AT). The increased power to identify treatment effects of automated scoring methodologies (MF and CP) supports the notion that human observer bias could be a major source of statistical power decrease in HKDT (Castaneda et al., 2012). Greater deviation between CP and visual estimates, however, suggests that this metric may be measuring a different biological response. Numerical differences between computational and visual scoring techniques is a known phenomenon in distance-based automated estimation of HKDT (Laursen et al., 2021; MacLean et al., 2022), although the deviation in these cases could originate from the sliding window approach used to summarize tracking data.

**Table 1.** Effect of temperature on mean estimates of thermal tolerance and concordance correlation coefficients between computational and visual datasets. Means not sharing any letter are significantly different by the Tukey-test at the 5% level of significance.

Critical thermal maxima
|  | *AT* | *CP* | *MF* | *Vis* |
| --- | --- | --- | --- | --- |
|  | Estimate | Estimate | Estimate | Estimate |
|  | (°C) | (°C) | (°C) | (°C) |
| 18°C | 39.6 a | 38.8 a | 39.5 a | 39.9 a |
| 25°C | 39.9 a | 38.7 a | 39.8 a | 40.2 a |
| 30°C | 40.7 b | 39.5 b | 40.6 b | 40.8 b |
| CCC | 0.65 | 0.06 | 0.61 |  |
| *Cb* | 0.96 | 0.46 | 0.91 |  |
| $\rho$ | 0.68 | 0.14 | 0.67 | |

Critical thermal minima
|  | *AT* | *MF* | *Vis* |
| --- | --- | --- | --- |
|  | Estimate | Estimate | Estimate |
|  | (°C) | (°C) | (°C) |
| 18°C | 3.25 a | 4.11 a | 3.19 a |
| 25°C | 5.85 b | 5.93 b | 5.91 b |
| 30°C | 7.12 c | 7.25 c | 7.33 c |
| CCC | 0.85 | 0.68 |  |
| *Cb* | 0.99 | 0.99 |  |
| $\rho$ | 0.85 | 0.69 | |

Heat knock down time
|  | *AT* | *CP* | *MF* | *Vis* |
| --- | --- | --- | --- | --- |
|  | Estimate | Estimate | Estimate | Estimate |
|  | (min) | (min) | (min) | (min) |
| 18°C | 79.5 a | 61.1 a | 77.4 a | 81.4 a |
| 25°C | 75.1 a | 60.3 a | 74.3 a,b | 74.4 a |
| 30°C | 84.8 a | 72.5 b | 86.4 b | 85.9 a |
| CCC | 0.90 | 0.60 | 0.85 |  |
| *Cb* | 1.00 | 0.66 | 0.98 |  |
| $\rho$ | 0.90 | 0.91 | 0.86 | |

Chill coma recovery time
|  | *AT* | *MF* | *Vis* |
| --- | --- | --- | --- |
|  | Estimate | Estimate | Estimate |
|  | (min) | (min) | (min) |
| 18°C | 23.2 a | 20.9 a | 21.6 a |
| 25°C | 28.8 b | 24.4 a | 28.7 b |
| 30°C | 25.8 a,b | 20.7 a | 23.5 a |
| CCC | 0.50 | 0.60 |  |
| *Cb* | 0.98 | 0.94 |  |
| $\rho$ | 0.51 | 0.64 | |

**Figure 1.**
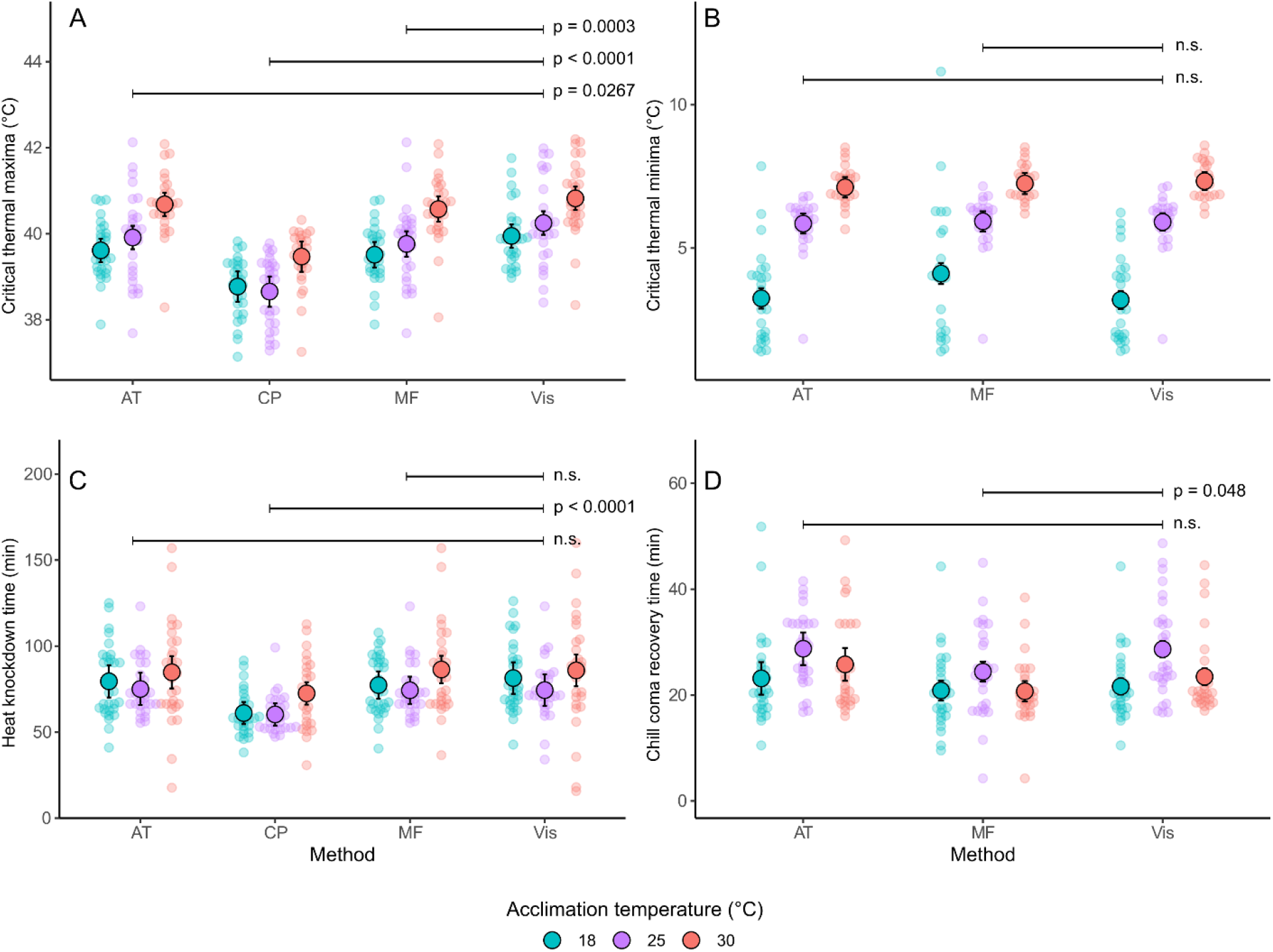
Average differences between computational and visual estimation of thermal tolerance in adult of *Drosophila melanogaster* flies: critical thermal maxima (A; n_AT_ = n_CP_ = n_MF_ = n_Vis_ = 90), critical thermal minima (B; n_AT_ = n_MF_ = n_Vis_ = 78), heat knockdown time (C; n_AT_ = n_MF_ = n_Vis_ = 89, n_CP_ = 87), and chill coma recovery time (D; n_AT_ = 85, n_MF_ = n_Vis_ = 88). Large black-border circles represent mean treatment values and bars represent s.e.m.; smaller circles depict raw data.

In most cases, the computational methods recapitulated the effects of acclimation on thermal endpoints. For both CTmax and CTmin, there was a clear acclimation effect present in all four scoring methods, with both CTmax and CTmin increasing monotonically as a function of acclimation temperature (Fig. 1). The acclimation effect for HKDT was less pronounced, but for all measures flies acclimated to 30°C had the highest knockdown time (Fig. 1). However, neither of the computational methods evaluated for CCRT (AT and MF) were unable to recover the treatment effects to the same degree of visual estimates. The reduced amount of motion data in CCRT could be biasing the threshold used in computational scoring, confounding background noise with motion in specific cases (see below).

### Source of disagreement between pairs of measurements

The reliability of computational methods to score thermal tolerance was assessed with the concordance correlation coefficient (CCC). To further assess the source of disagreement, CCC was decomposed into its accurateness (C_b_) and precision (ρ) components. The lowest CCC values were observed between CP and visual estimates, confirming our suspicion that this statistical interpretation of the data could be measuring an alternative component of thermal performance, albeit with high correlation to HKDT. Estimates obtained with AT and MF present high accuracy but variable precision across assays (Table 2), but together are either comparable (CTmax, CCRT) or even more agreeable (CTmin, HKDT) to visual estimates than previously reported inter- and intra-researcher consistency for HKDT (Castaneda et al., 2012). High accuracy present in heuristic-inspired (AT and MF) computational methods confirms their capacity to reliably measure the mean and variance of thermal tolerance endpoints. Moderate precision suggest that average differences are influenced by discrepancies between individual pairs of measurements possibly originating from an interaction between biological and technical sources of error. We identified pairs of estimates with outlying differences between visual and MF datasets using a graphical technique (Fig. 2A-D). Experimental subjects with discrepant scores were identified outside the 95% limits of agreement (Bland and Altman, 1986). Within the limits of agreement, the dispersion of the differences in lower thermal limits (CTmin and CCRT) was closer to zero (Fig. 2B,D) while greater dispersion was observed in upper thermal limits (CT_max_ and HKDT) owing to the greater amount of motion information available when experimental subjects are exposed to heat (Fig. 2A,C). Outside the limits of agreement, disagreeable individuals were computationally underestimated in CT_max_ (11 out of 12), CT_min_ (6 out of 7), and CCRT (7 out of 8) but overestimated in HKDT (5 out of 6).

**Figure 2.**
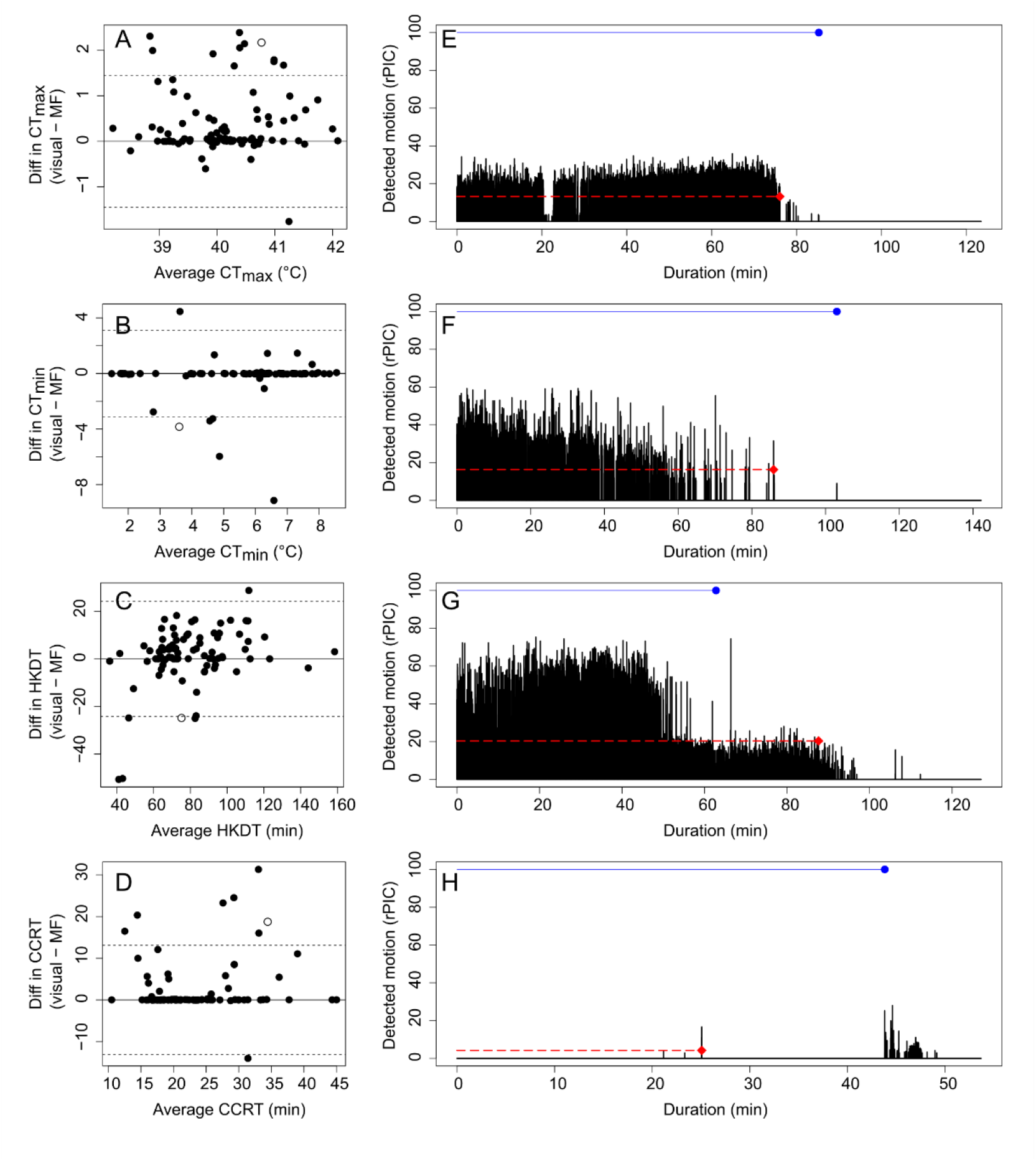
Agreement test between Visual and Median Frame estimates for CT_max_ (A, n = 90), CT_min_ (B, n = 78), HKDT (C, n = 89), and CCRT (D, n = 88) and a motion plots (E-H) of the disagreeable individuals (empty circles in A-D). Length of the red long-dashed line indicates the time the individual was active, or inactive in the case of CCRT, height indicates the threshold used to score the thermal limit using the MF method, and the red diamond at its end indicates the endpoint. The visual estimate is illustrated with a blue solid line at 100 rPIC.

Finally, we investigated the source disagreement in some of these cases with individual plots of motion (Fig. 2E-H). Motion plots of outlying individuals (Fig. 2E-H corresponding to empty circles in Fig. 2A-D) suggest that the source of disparity originates from the threshold value (height of long-dashed red line). In dynamic assays, the threshold was too high (low sensitivity) to detect the motion events used to score the endpoint visually (Fig. 2E-F), but the effect was inverse in HKDT outliers (Fig. 2G) that scored background noise as motion due to a lower threshold value (high sensitivity). In a similar manner, the threshold used to score the CCRT outlier (Fig. 2H) provided an earlier recovery time by identifying background noise (i.e., an autofocus artifact) as motion. Nonetheless, discrepancies can be identified and reassessed with a visual inspection of the motion plots.

Our results indicate that automated methods of endpoint estimation (AT and MF) are appropriate substitutes for visually obtained estimates of knockdown-based assays (CT_max_, CT_min_, and HDKT). Our results also indicate that the MF method is preferred, as it consistently recapitulated biological treatment effects and does not require calibration. Still, these methods present limitations when scoring recovery from chill coma, perhaps because the sparse motion data in this type of assay that does not inform of a consistent threshold between active and inactive states. Additionally, our motion detection strategy with DIME provides the opportunity to interpret motor activity as a quantitative expression with the use of motion plots. Taken together, our results indicate that automated methods for scoring thermal tolerance can provide comparable values to classic methods relying on manual scoring.

## Acknowledgements

We would like to acknowledge Ioulia Bespalova, Luis E. Castañeda, and Juan Soto Hernández for their feedback on previous versions of the application DIME.

## Data availability

The command-line application DIME can be downloaded from https://github.com/fernan9/DIME

